# Proton uptake mechanism in bacteriorhodopsin captured by serial synchrotron crystallography

**DOI:** 10.1101/576629

**Authors:** Tobias Weinert, Petr Skopintsev, Daniel James, Florian Dworkowski, Ezequiel Panepucci, Demet Kekilli, Antonia Furrer, Steffen Brünle, Sandra Mous, Dmitry Ozerov, Przemyslaw Nogly, Meitian Wang, Jörg Standfuss

## Abstract

Conformational dynamics are essential for proteins to function. Here we describe how we adapted time-resolved serial crystallography developed at X-ray lasers to visualize protein motions using synchrotrons. We recorded the structural changes upon proton pumping in bacteriorhodopsin over 200 ms in time. The snapshot from the first 5 ms after photoactivation shows structural changes associated with proton release at comparable quality to previous X-ray laser experiments. From 10-15 ms onwards we observe large additional structural rearrangements up to 9 Å on the cytoplasmic side. Rotation of Leu93 and Phe219 opens a hydrophobic barrier leading to the formation of a water chain connecting the intracellular Asp96 with the retinal Schiff base. The formation of this proton wire recharges the membrane pump with a proton for the next cycle.

## Main Text

Proteins are intrinsically dynamic molecules that follow defined conformational rearrangements to drive the fundamental biochemistry of life. Structural studies of these dynamic processes are challenging, because structural intermediates decay and are only reliable in wild-type proteins at physiological temperatures. Pump-probe crystallography offers the unique opportunity to capture successive snapshots of light-activated processes to study structural changes at the level of individual atoms (Neutze & Moffat 2012).

X-ray free electron lasers (XFELs) and the emerging method of time-resolved serial femtosecond crystallography (TR-SFX) brought exciting new opportunities for structural biologists (Schlichting 2015; Chapman 2019). Recent studies on the photoactive yellow protein (Tenboer et al. 2014; Pande et al. 2016) proved that TR-SFX is a powerful new tool to study the structural dynamics of proteins. The ultrafast but bright X-ray pulses from these powerful linear accelerators outrun radiation damage and allow the use of small crystals, resulting in high protein activation levels. The serial injection of crystals further favors the study of non-reversible reactions and the collection of redundant datasets of high quality. The great potential of the method to study membrane transport was realized when a whole series of structural states of the light-driven proton pump bacteriorhodopsin (bR), ranging from the femtosecond to the early millisecond regime, were determined by TR-SFX.

Primary energy conversion processes in biology are variations around one central theme: proton transfer across membranes. Time-resolved crystallographic studies provided much insight into how proton pumping is achieved in bR. In particular they focused on the early events in the pumping cycle including the ultrafast process of energy capture through photoisomerization of retinal (Nogly et al. 2018) and the proton release step from the retinal Schiff base (SB) towards the extracellular side of the membrane (Nango et al. 2016). The fundamental piece missing in our dynamic view of proton transport is how the protein rearranges to reprotonate the SB from the cytoplasmic side to reload substrate for the next pumping cycle.

We answer this question by adapting injector-based serial crystallography for time-resolved measurements in the millisecond range at widely available synchrotron sources. Requirements for time-resolved crystallography using the Laue method were never met for bR (Wickstrand et al. n.d.). Therefore, pioneering works to determine the structures of spectroscopic intermediates were carried out using freeze-trapping and mutagenesis (Wickstrand et al. 2015). Yet major controversies remained because mutations modify critical activation switches and structures of mutants may not adequately reflect rearrangements in the native protein. Such structures may further suffer from radiation damage, are obtained at non-physiological temperatures and lack the temporal dimension critical to understanding protein dynamics. Serial millisecond crystallography (SMX) approaches work well with high viscosity injectors (Weierstall et al. 2014), as the injectors extrude crystals slowly and allow sufficient X-ray exposure to collect high-resolution data with the photon flux available at synchrotrons (Grünbein & Nass Kovacs 2019). Our recent work has shown that modern high frame-rate and low-noise detectors make SMX a viable method for routine room-temperature structure determination (Weinert et al. 2017). By extending the setup with a simple class 3R laser diode we enable time-resolved studies with millisecond time resolution (TR-SMX) and bring dynamic serial crystallography from the XFEL niche to the large community of synchrotron users worldwide. We demonstrate the technology by recording the temporal evolution of the two structurally distinct M and N intermediate states of bR and how they decay back to the dark-state. The M-state structure obtained by TR-SMX shows the structural rearrangements necessary for proton release with comparable quality and in agreement with previous XFEL experiments. The N-state is characterized by large conformational rearrangements up to 9 Å on the cytoplasmic side not yet resolved by dynamic crystallography. These open the protein for the formation of a Grotthuss proton wire, a transient water chain that allows the reprotonation of the Schiff base (SB) from Asp96 the primary proton donor at the cytoplasmic side of the membrane. The observations align with the classical alternate access model of membrane transport and further complete the dynamic view of vectorial proton transport in bR by resolving the critical proton uptake step at the end of the pumping cycle.

## Time-resolved serial millisecond crystallography

The most straightforward experiment possible with our TR-SMX setup (Figure 1) leaves the laser diode on while data from continuously light-activated protein molecules within the crystal are being collected. By comparing these data to serial data collected without illumination (**Figure S1**) it is possible to reveal dominant conformational changes in a photocycle analogous to steady-state spectroscopic techniques. From bR microcrystals with a size of about 35×35×3 μm^3^, we collected 114’052 light and 119’374 dark diffraction patterns in the steady-state mode with data extending to a resolution of 1.8 Å (**Table S1**). The quality of difference electron density maps (F_o_(light)-F_o_(dark)) improved with the number of included images with characteristic difference density peaks starting to plateau at about 10’000 images, indicating data convergence (**Figure S2**). The estimated excitation level was 38% and resulted in well-resolved difference density map features, with the strongest positive and negative peaks at 7.3 σ and 8.5 σ, respectively. Structure refinement against the steady-state data (**Table S1**) revealed two activated species of bR (**Table S2**). While one subspecies had been observed in previous TR-SFX structures obtained at 1.7 ms and 8.3 ms after photoactivation, the second subspecies featured large rearrangements on the cytoplasmatic side indicating that the dynamic view of proton pumping in bR had not yet been completed.

**Figure 1.**
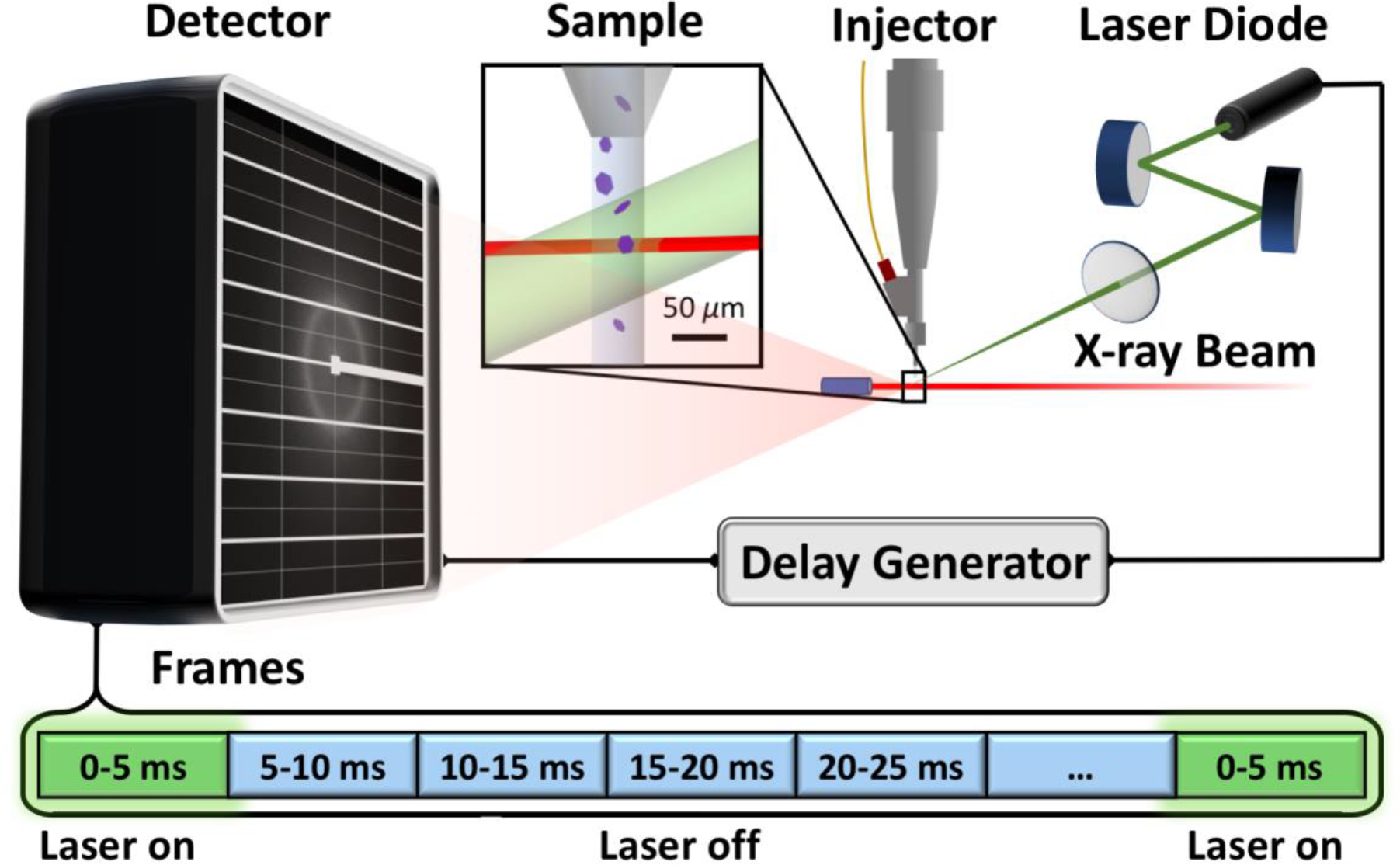
Experimental pump-probe setup and TR-SMX data collection. The laser (green) originates from a class 3R laser diode and is controlled via two steering mirrors before being focused to intersect with the extrusion path of the high viscosity injector (grey) and the X-rays (red). The laser and detector are triggered by a delay generator. Running the detector at 200 Hz, the data collection cycle consists of 40 frames each 5 ms long. The laser is activated for 5 ms during collection of the first data frame and is then turned off for the remaining frames during which the illuminated bR completes its photocycle.

In order to follow the rise and decay of these intermediates and to obtain separate structures, we developed a data collection scheme, which triggers the laser diode using a transistor-transistor-logic signal issued from a delay generator to achieve laser exposure synchronized with data collection at the detector (Figure 1). In this setup bR crystals were exposed for 5 ms with laser light at the X-ray interaction region, while tracking the photoreaction by collecting 5 ms long timepoints over 40 consecutive frames. This data collection sequence was continuously repeated while new crystals were resupplied by the high viscosity injector (Weierstall et al. 2014). After about 5 hours of data collection, each of the timepoints contained ~13’000 indexable diffraction patterns with the resolution extending to 2.3 Å (**Table S1**).

Analysis of the data provides a ‘continuous’ molecular movie of structural changes over 200 ms in time during which bR molecules complete their photocycle within the crystal. Within this time frame we observe a long-term decay of difference map signal and complete restoration of the dark-state within about 100 ms (**Movie S1**). The rate of this long-term decay (**Figure S3**) and the evolution of structural states as determined by occupancy refinements (**Figure S2**) occur within the temporal range established by time-resolved spectroscopy on bR crystals(Nogly et al. 2016; Nango et al. 2016; Efremov et al. 2006).

These results demonstrate that time-resolved serial crystallography is not limited to the few XFELs worldwide but can also be done efficiently at widely available synchrotrons. Results are of comparable quality to our previous time-resolved studies at XFELs on the same crystals (**Figure S4**), even though the resolution was limited by the available photon flux. However, the resolution is compensated by high levels of activation from the millisecond long light exposure, more accurate data due to the use of a photon counting detector and a more stable monochromatic beam as previously observed during *de novo* phasing (Weinert et al. 2017). Due to the slow extrusion rate, less than 20 µl of prepared microcrystals were necessary to obtain these data instead of milliliters needed for the XFEL experiments. The reduced sample consumption significantly lowers the entry barrier for time-resolved crystallographic studies. While the simple TR-SMX setup implemented at the Swiss Light Source delivers excellent results, there is great potential to improve the method further. Pink beams (Meents et al. 2017) and diffraction limited sources (Eriksson et al. 2014) optimize beam characteristics for faster data acquisition. The Hadamard transform method (Yorke et al. 2014) and continued software development (White et al. 2016) are means to improve data analysis. Together with stronger nanosecond lasers as well as new fast and accurate detectors (Leonarski et al. 2018) these improvements will allow the time resolution to be pushed from milliseconds into the lower microsecond regime to cover the majority of biologically relevant protein dynamics.

## Completing the bR pumping cycle

The molecular movies of structural changes in bR are an impressive example for how XFEL sources can be used to understand biology (Wickstrand et al. n.d.). Our TR-SMX measurements extend the dynamic view of bR into the second half of the photocycle (Figure 3), where the proton substrate is reloaded.

Structures refined against the 0-5 ms timepoint (**Table S2**) were a good fit to M-state structures obtained by cryo-trapping (Takeda et al. 2004) or TR-SFX (Nango et al. 2016; Nogly et al. 2018) (r.m.s.d. 0.463 Å for the cryo-trapped structure 1IW9, 0.397 Å for 1.7 ms structure 5B6Z and 0.269 for the 8.3 ms structure 6G7L). Difference maps obtained from 10-15 ms onwards however showed clear additional features at the cytoplasmic side of the protein (Figure 2). We found a better fit in the dark-state X-ray (Wang et al. 2013) and electron diffraction structures (Subramanlam & Henderson 2000) of the constitutively open Asp96Gly/Phe171Cys/Phe219Leu triple mutant (r.m.s.d. of 0.586 Å for the X-ray structure 4FPD and 0.845 Å for the electron diffraction structure 1FBK), which are considered analogous to the wild-type N-state obtained by photoactivation. The structural rearrangements in the N-state with respect to the dark and M-states occur in the cytoplasmic ends of helices E, F, G. The largest change in helix F corresponds to a movement of 9 Å measured at the Cα position of Glu166 (**Movie S2**) and is consistent with transient movements of helices F and G in the millisecond range observed by time-resolved spin labelling studies in purple membranes (Radzwill et al. 2001). The final O-state does not accumulate well in bR crystals (Efremov et al. 2006) and is in a fast equilibrium with the N-state (Souvignier & Gerwert 1992), suggesting that no large structural rearrangements occur in the transition. The small changes upon relaxation of retinal from the *cis*-isomer in the O-state back to the *trans*-isomer (Zhang et al. 2012) are spread out in time which makes them difficult to detect. Our TR-SMX experiment thus completes the crystallographic view of the bR photocycle, by revealing the rearrangements from the M-to the N-state and back to the dark state.

**Figure 2.**
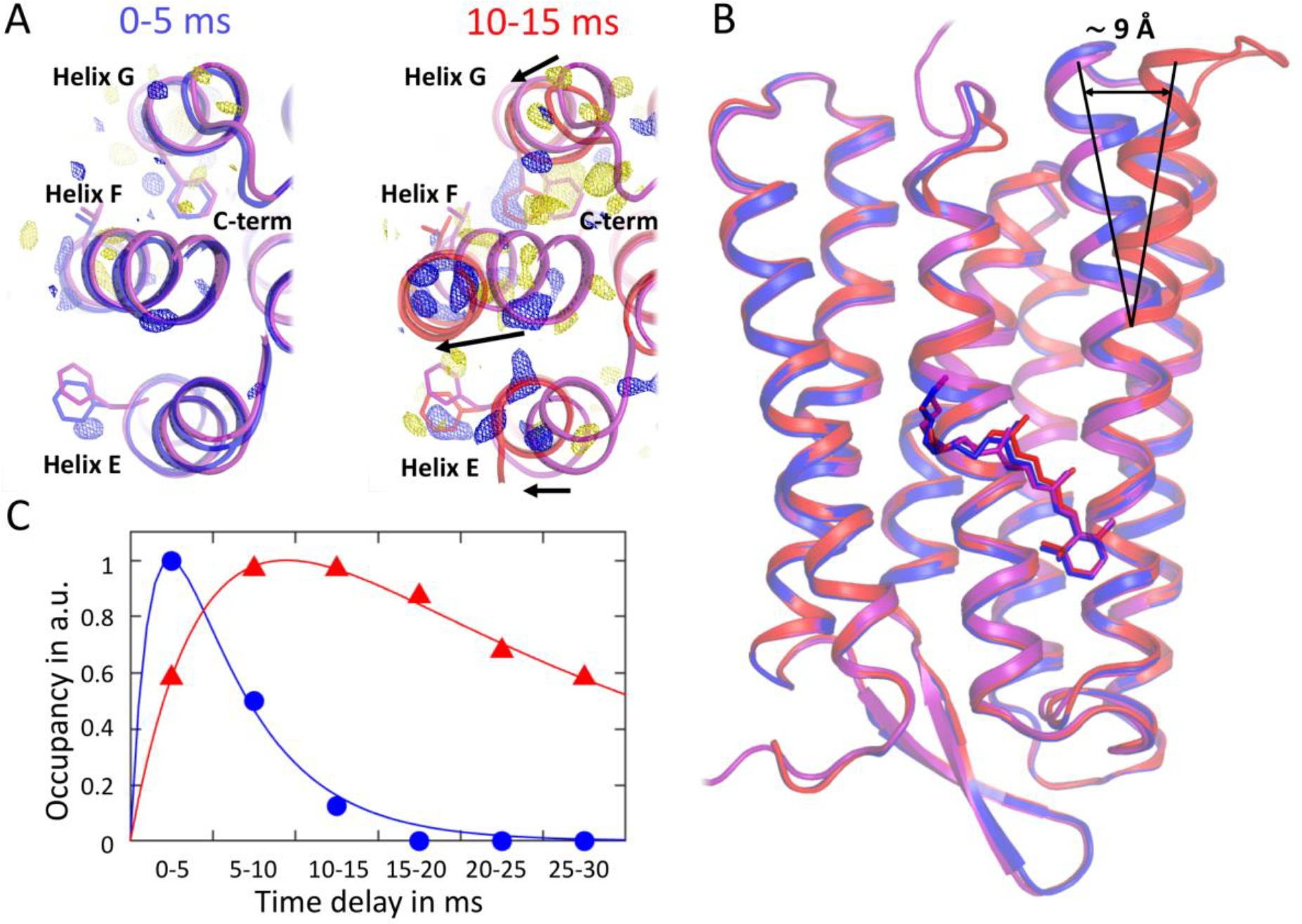
Evolution of structural intermediates over time. (**A**) Large scale conformational changes on the cytoplasmic ends of helices E, F and G (dark-state in purple, M-state at 0-5ms in blue and N-state at 10-15ms in red) are evident from difference electron density maps (left: F_o_(0-5ms)-F_o_(dark), right: F_o_(10-15ms)-F_o_(dark) at 3σ with positive difference density in blue and negative difference density in yellow). (**B**) The side view illustrates the large rearrangements on the cytoplasmic side of ~9 Å at the end of helix F (black lines). (**C**) Occupancy refinement of the different states (M-state in blue circles, N-state in red triangles) allows following the structural evolution of the two intermediates over time. Biexponential fits against the normalized data are shown with solid lines.

Structural changes in helices E and F involve large side chain motions. Most prominently, the aromatic ring of Phe156 moves 9 Å and replaces the ring of Phe171 to alter the hydrophobic interaction network holding helix F in its original position. Moving helix F goes hand in hand with the disordering of the C-terminus starting from Arg227 at the end of helix G (Figure 4). In contrast, the structural changes in helix C are dominated by smaller sidechain rearrangements. The largest conformational change in helix C is a rotamer change of Leu93 which goes hand in hand with a displacement of the nearby Phe219 sidechain in helix G (**Movie S2**). These changes create space for the formation of a water molecule chain (Wat404, Wat453, Wat454) observed as strong positive difference map peaks (F_o_(10-15ms)-F_o_(dark) above 3σ) connecting Asp96 on the cytoplasmic side with the retinal SB (Figure 3). Additional experimental evidence for such waters originates from a combination of Fourier transform infrared spectroscopy, site directed mutagenesis and molecular dynamic simulations (Gerwert et al. 1989; Freier et al. 2011). Our experimental results are consistent with molecular dynamic simulations showing that three waters are sufficient to close the gap and enable the proton transfer step to the SB (Freier et al. 2011). The positions of the three water molecules are different from the four waters observed in a freeze trapping structure of the Val49Ala mutant (Schobert et al. 2003) and the state trapped by mutagenesis lacks the large conformational changes on the cytoplasmic site. The constitutively open triple mutant Asp96Gly/Phe171Cys/Phe219Leu on the other hand allowed to trap N-like features without light activation (Wang et al. 2013). However, removing the charge from Asp96 and reducing the size of the Phe219 side chain prevented formation of the water chain (**Figure S5**). Such controversies have long hindered the emergence of a consensus concerning large conformational changes in the late bR photocycle (Wickstrand et al. 2015). The useful mutations by their very nature had to target functionally critical regions which complicated interpretation. Our study resolves these discrepancies by showing the relevant arrangements in light-activated native bR measured at physiological temperatures and with real-time resolution.

**Figure 3.**
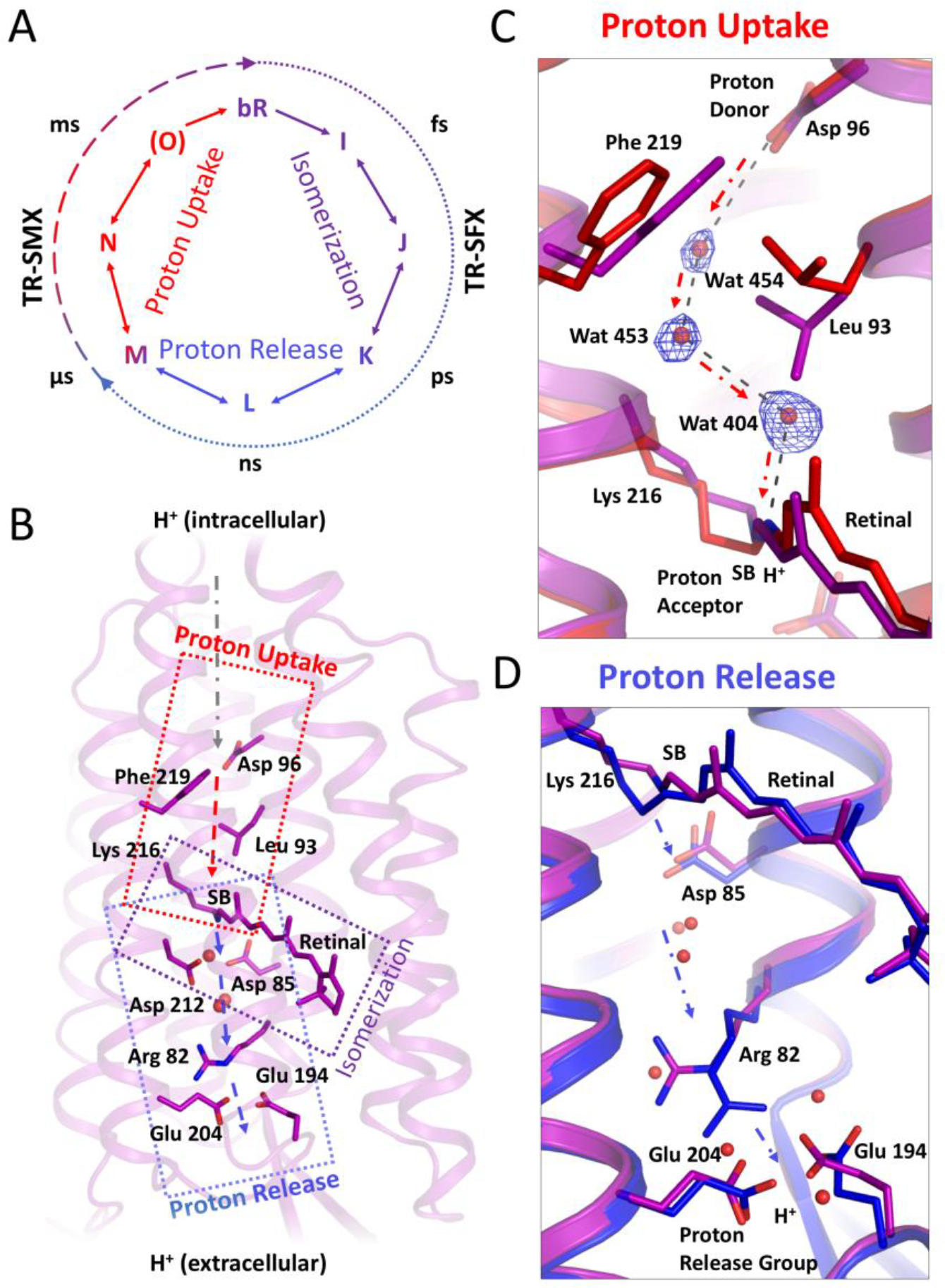
Proton pumping in bacteriorhodopsin proceeds via three principal steps. **(A)** The proton pumping cycle in bR can be followed through a series of spectroscopic intermediates (I, J, K, L, M, N, O) occurring within a few hundred femtoseconds up to tens of milliseconds after activation. **(B)** The first principal step is the light-induced *trans-cis* isomerization of the retinal chromophore and deprotonation of the SB link to the protein. The following conformational changes allow transfer of the proton along a series of key amino acid residues (sticks) and water molecules (spheres) to the release group at the extracellular side of the membrane. In the last principal step, the SB is recharged with a proton from the intracellular side. (**C**) At 10-15 ms we observe the opening of the hydrophobic barrier between the primary proton donor Asp96 to the SB (dark-state in purple, N-state refined with 10-15 ms data in red). The changes are accompanied by the ordering of three water molecules (positive F_o_(10-15ms) - F_o_(dark) density in blue at 3 σ). (**D**) The 0-5 ms timepoint shows the rearrangements necessary for proton release in agreement with previous XFEL studies (Nango et al. 2016; Nogly et al. 2018) (dark-state in purple, M-state refined with 0-5ms data in blue). Arrows indicate the direction of proton transfer reactions.

**Figure 4.**
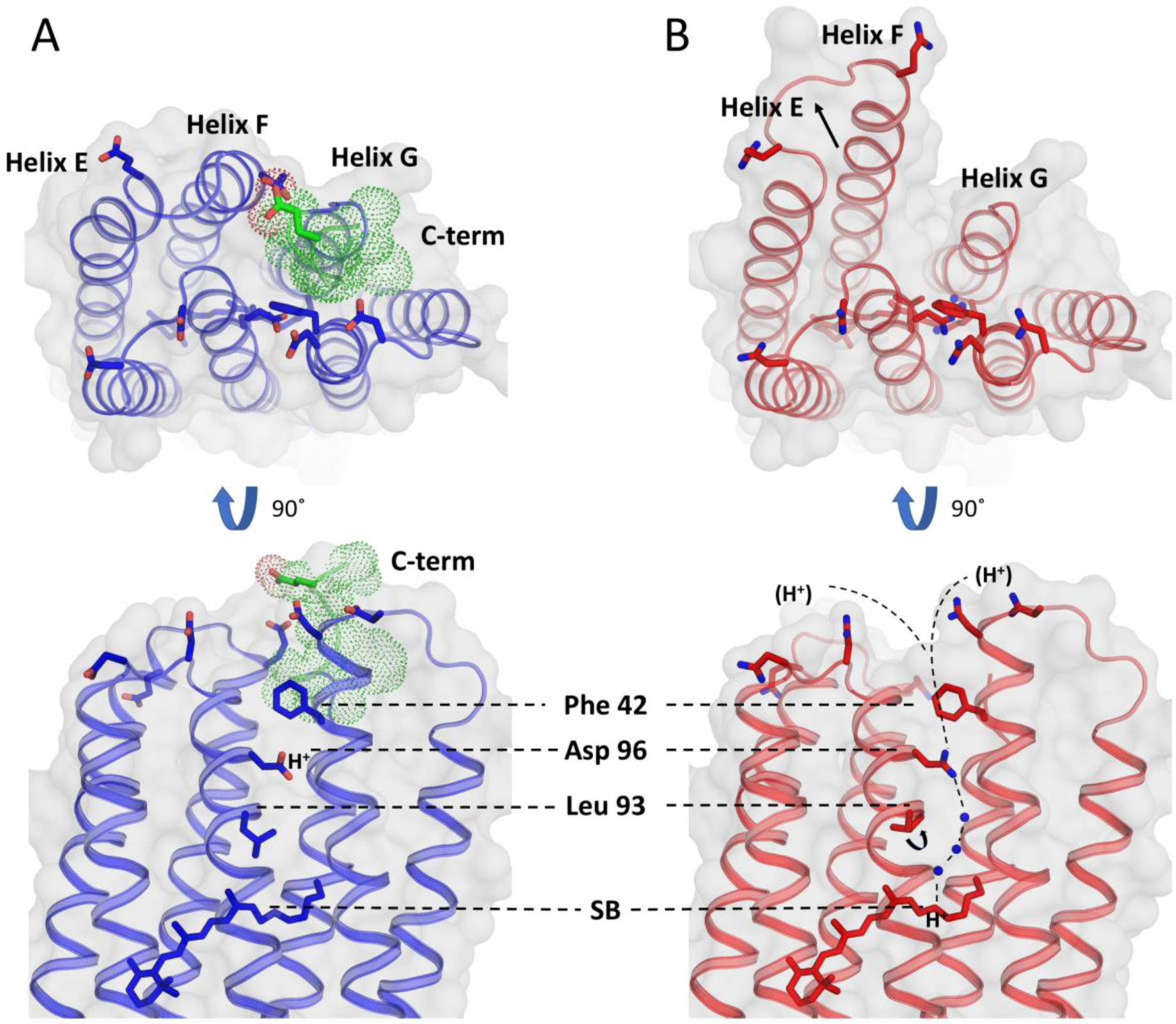
Opening of the cytoplasmic side facilitates reprotonation. Rearrangements of helices E, F and G between the M-state with a protonated Asp96 **(A)** and the N-state with a protonated SB **(B)** open the cytoplasmic side (protein surface shown in grey) and lead to disordering of the C-terminus (green dotted spheres). The proton position is indicated on either Asp96 or the SB and the connecting proton wire forming in the N-state is highlighted. The structural rearrangements change the chemical environment of charged residues on the cytoplasmic surface (including Glu232, Glu166, Glu161, Asp102, Asp104, Asp38, Asp36 shown as sticks) that have been implicated in funneling protons to reprotonate Asp96 (reprotonation paths indicated as dashed lines).

## Molecular mechanism of proton uptake

The structural transition fits well with the molecular mechanism of vectorial proton pumping (**Movie S2**) (Gerwert et al. 2014). Mechanistically, bR can be divided into an extracellular half and a cytoplasmic half with retinal positioned at the center (Figure 3). Light-induced isomerization of the retinal chromophore is the first principal step in the reaction and provides the energy for proton pumping. The energy is used in the second principal step to drive protein conformational changes on the extracellular side to transfer a proton from the retinal SB towards the extracellular release group via a series of residues and water molecules. Transferring the proton to the release group removes the substrate from the center of the membrane.

In the third principal step, the protein has to adopt a different conformation to allow the SB to accept a proton from the intracellular side of the membrane. Preventing proton back leakage to the cytoplasmic side is critical for vectorial transport against a concentration gradient. In bR, this is achieved through a hydrophobic barrier between the primary acceptor of protons in the center (the SB) and the proton donor on the cytoplasmic side of the membrane (Asp96). This 12 Å barrier has to be opened at the right time during each cycle to allow the SB to accept a new proton and recharge the system. The formation of a water chain, oriented by the protein environment, could principally act as a Grotthuss proton wire (Agmon 1995), bridging the hydrophobic barrier to deliver a new proton to the SB. In the 10-15 ms timepoint, the rotation of Leu93 together with that of Phe219 provides the space for three water molecules that connect the SB with the intracellular proton donor Asp96 (Figure 3), opening the hydrophobic barrier. Transferring the proton to the SB replenishes the substrate at the center of the membrane.

The structural rearrangements we observe leave Asp96 covered by a single hydrophobic lid formed by Phe42, making it accessible for reprotonation. Furthermore, structural changes to a cluster of residues acting as a proton funnel at the cytoplasmic side of the protein aid reprotonation of Asp96 (**Figure4**) (Riesle et al. 1996), which may transiently proceed via an intracellular uptake cluster (Lorenz-Fonfria et al. 2017).

Time-resolved serial crystallography revealed the structural reorganizations during proton uptake and release in bR with astounding detail and over many orders of magnitude in time. The alternating rearrangements on the extracellular and intracellular sides during the pumping cycle of bR agree with an alternate access model and may provide a template to understand the principal transport steps in other membrane transport proteins. The large-scale motion of helix F further resembles the outward movement of transmembrane helix 6 in G protein-coupled receptors. Bringing time-resolved serial crystallography developed at XFELs to the broader scientific community using synchrotrons will answer many other pressing questions in structural biology.

## Materials and Methods

### Lipidic cubic phase crystallization and sample preparation

The purification and crystallization of bacteriorhodopsin were performed as described previously (Nogly et al. 2018). Purple membranes from *Halobacterium salinarum* were solubilized overnight in the presence of 1.2% b-octylglucoside (Anatrace) and 50 mM sodium phosphate buffer pH 6.9 (GERBU). After solubilization the pH was adjusted to 5.5 with 0.1 M HCl and the insoluble fraction was removed by 1 hour of centrifugation at 150’000 × g. Before crystallization the protein was concentrated to 40 to 80 mg ml^−1^ using Millipore centricolumns with a 50 kDa cutoff. Lipidic Cubic Phase (LCP) for crystallization was obtained in a Hamilton syringe by mixing protein with monoolein (Nu-Chek) in a 42:58 ratio. Subsequently the LCP was slowly injected into another Hamilton syringe filled with crystallization buffer consisting of 100 mM Sorensen buffer pH 5.6 (GERBU) and 30% polyethylene glycol 2’000 (Molecular Dimensions). The crystallization was carried out at 21°C and crystals with an approximate average size of 35×35×3 µm^3^ were obtained within 3 to 6 days. All purification and crystallization steps were performed under dim red light or in the dark.

Shortly before the SMX experiment, the crystallization buffer was removed from LCP containing crystals via a syringe coupler by slow pressing on the syringe plunger. Crystals were incubated in 200 mM Sorensen buffer overnight to wash out polyethylene glycol and favor formation of LCP. Further monoolein was added to bring the sample into a homogeneous and transparent LCP mesophase. The crystal density within the sample was homogenized using a custom made “3-way syringe coupler” (James et al. 2018) before loading the high viscosity injector.

### Laser setup

The simple Class 3R diode laser we integrated into our SMX setup (Weinert et al. 2017) is inexpensive, available at a range of wavelengths and requires significantly less strict safety protocols than the Class 3B and 4 lasers typically used for time-resolved experiments. The laser diode can be triggered at up to 10 kHz and provides sufficient light to “pump” each chromophore many times within a millisecond exposure to allow high excitation levels.

The laser delivery system consists of 4 mirrors (Thorlabs), a 100 mm focal length achromatic objective lens (Thorlabs), and a laser diode module (Roithner Lasertechnik). The system is placed on an optical breadboard custom fitted to the beamline to allow for optical components to be mounted level with the sample position. The entire system is rigidly mounted in a 30 mm optical cage system (Thorlabs), which allows for rough beam alignment by moving the position of the cage by hand. The final two mirrors in the system allow for fine beam positioning on the sample. Beam focus is achieved by sliding the objective lens along the cage rails for a coarse focus, and fine focus is provided by a z-axis translatable lens mount. The laser diode module emits a 5 mW, 3 mm diameter collimated beam at 520 nm. The resultant beam spot dimensions after focusing (104 µm by 173 µm 1/e^2^) were measured by a knife edge scan. The laser power was measured as 2.84 mW at the position where the LCP extrusion crosses the X-ray path which translates into a maximal power density of 37.43 W/cm^2^ in each 5 ms laser exposure.

### Data collection

Data for steady-state SMX were collected by interleaving 30 minute intervals during which we switched the laser on or off (**Supplementary Figure 1**). The crystal-laden LCP was injected in the intersecting X-ray and laser beam paths by a high viscosity injector (Weierstall et al. 2014) with a 50 μm diameter nozzle. The jet speed was 250 μm/s and data frames were recorded using an Eiger16M detector with a frame rate of 50 Hz. Together with a 20 × 5 μm beam this resulted in an extrusion by one beam height (5 μm) during the collection of one frame, minimizing similarities in consecutive frames for more reliable statistics.

For TR-SMX experiments the jet speed was also 250 μm/s and data frames were recorded using an Eiger16M detector in a 4M region-of-interest mode with a frame rate of 200 Hz. Together with a 20 × 5 μm beam this resulted in a translation of the jet by a quarter of a beam height (1.25 μm) during the collection of one frame, maximizing similarities in consecutive time points for more comparable difference maps. A TR-SMX data collection cycle consisted of 40 frames each 5 ms long. The laser diode was activated for 5 ms during collection of the first data frame and then turned off during the collection of the remaining frames during which the illuminated bR completed its photocycle (Figure 1). This scheme allowed a continuous data collection with enough time for the activated fraction of bR molecules to decay to the dark-state (**Supplementary Figure 4**). Control Difference density maps between all dark data and the last 5 frames from the TR-SMX data collections were flat (**Supplementary Movie 1**). For maximal consistency, we therefore used these last 5 frames as dark data for the calculation of TR-SMX difference maps and extrapolated maps in the TR-SMX experiment (**TR-SMX dark, Supplementary Table 1**).

The photon flux of 1.5 × 10^12^ photons/s resulted in a radiation dose of about 5 kGy per recorded detector frame (at 200 Hz) when calculated with RADDOSE-3D (Zeldin et al. 2013), resulting in 5 to 140 kGy per crystal depending on their size and orientation while traversing the beam.

### Data processing

All data were indexed, integrated and merged using Crystfel (White et al. 2012; White et al. 2016) version 0.7.0. Data were indexed using the mosflm and dirax algorithms. Data were integrated using the --rings option in indexamajig. Patterns were merged using partialator with the following options: --model=unity, --iterations=1, --push-res=1.5.

### Difference map calculation

F_o_(light)-F_o_(dark) difference maps were calculated using PHENIX (Adams et al. 2010). All F_o_(light)-F_o_(dark) difference maps were calculated excluding amplitudes smaller than 3 σ and resolutions lower than 10 Å and using the multi-scaling option. All difference maps for comparing TR-SMX with TR-SFX were calculated with the same high-resolution threshold of 2.1 Å using data from (Nango et al. 2016) (1.7 ms SACLA) and (Nogly et al. 2018) (8.3 ms LCLS).

### Model building and refinement

All structural refinements were done using PHENIX (Adams et al. 2010) with iterative cycles of manual adjustments made in Coot (Emsley & Cowtan 2004).

### Steady-state SMX

The SFX dark-state model (PDB code 6G7H (Nogly et al. 2018)) was directly refined against the light off data (Steady-SMX dark, **Supplementary Table 1**). Extrapolated data were calculated as described elsewhere (Nogly et al. 2018). The activation level was determined by calculating extrapolated maps using the steady-SMX light data and the steady-SMX dark data at different activation levels in steps of 2 % until features of the dark-state appeared at the retinal, Arg 82 and in helices E, F and G. Based on this analysis an activation level of 38 % was chosen. Negative amplitudes resulting from the extrapolation procedure were removed and sigmas of steady-SMX light data were used in refinement against extrapolated data. Initially, the structural model of the M-State determined by TR-SFX (5B7Z) was chosen for refinement. The activated state included in 5B7Z was superposed on our refined steady-SMX dark model and then directly refined against extrapolated data. Further analysis using F_o_(steady-SMX light)-F_o_(steady-SMX dark) difference maps together with the extrapolated maps revealed that the M-state model was not sufficient to explain the data. Superposing of several structural models of bR showed the Asp96Gly/Phe171Cys/Phe219Leu triple mutant to be the best fit to the observed structural changes. Hence, we included the model in the refinement as an alternative conformation. The sequence of the model was modified back to the wild-type and the model was manually adjusted to best fit observed difference map features as well as extrapolated maps. Initially, the occupancy of both models was set to 50% and xyz coordinates as well as B-factors were refined. In later stages the occupancy was further refined resulting in an occupancy of 60% M-state and 40% N-state. In a final step, the water molecules identified in the time-resolved maps (Wat 404, 453, 554) were added based on difference electron density maps. The coordinates of water molecules identified based on time resolved maps were not refined.

### Time-resolved SMX

The F_o_(175-200 ms)-F_o_(Steady-SMX dark) difference map showed that no light features remained and the decay was complete to the detection limit; hence the 175-200 ms snapshot is termed TR-SMX dark. Extrapolated data and activation levels (**see Supplementary Table 2**) were calculated as described above for the steady-SMX data, but using the TR-SMX dark dataset and selected TR-SMX timepoints as light data. Negative amplitudes resulting from the extrapolation procedure were removed and sigmas of TR-SMX light data were used in refinement against extrapolated data. Structures representing the dominant state in the 0-5 ms and 10-15 ms timepoints were refined using the steady-state M-or N-states as starting models. In each case only the dominant conformation was chosen based on difference maps features and occupancy refinement.

### Occupancy refinement

The steady-state SMX dark and M/N models were combined into a single file with three alternative conformations. The model was refined against steady-state light data to obtain equilibrated starting occupancies for the three states. The B-factors of the model where then reset to the average value of 53 Å^2^ and only the occupancy was refined in Phenix over 10 cycles against data from seven timepoints (TR-SMX dark, 0-5 ms, 5-10 ms, 10-15 ms, 15-20 ms, 20-25 ms, 25-30 ms). The refined occupancies were then read out from the resulting PDB files. Correcting for dark-state contribution and normalizing the resulting occupancies to the largest value allows tracing the evolution of M and N-state over time (Figure 2C).

The rise and decay of the M and N intermediate states was further analyzed in Matlab using a global least-squares fit with biexponential function (*f* = ε ⋅[exp(−*t*/τ_*d*_) − exp(−*t*/τ_*r*_)] where *t* is time, τ_*d*_ and τ_*r*_ are time constants of state decay and rise, respectively, and *ε* is the effective protein excitation level. For each state, the decay time constant τ_*d*_ was set equal to rise time constant τ_*r*_ of a consequent state. The fit yielded a τ_*d*_ of 23.8 ± 2.7 ms for the N intermediate, which is in the range of the previously reported time constant for the crystalline bR N-state determined by spectroscopy (Efremov et al. 2006). The resulting M intermediate decay time constant was 4.7 ± 0.3 ms, and the fitted effective protein excitation was *ε* 19% ± 2% close to the experimentally determined level of 24% (compare sections on refinement above). The errors correspond to 95% confidence interval derived from the global fitting procedure.

## Acknowledgments

We thank Dieter Oesterhelt and Gebhard Schertler for providing purple membranes.

## Author contributions

T.W., P.S., M.W. and J.S. conceived the research with suggestions on laser setups from D.J. and F.D.. Protein and microcrystals were prepared by A.F. and P.N.. Slow crystal extrusion for the use at synchrotrons was optimized by P.S. and D.J.. Synchronized diode and detector triggering was implemented by E.P.. The pump-probe setup was built by D.J. and F.D. with suggestions from M.W.. Data collection was done by T.W., P.S., D.J., A.F., D.K., S.B., F.D., D.O., P.N., M.W. and J.S.. Data processing and structural refinements were done by T.W. The manuscript was written by T.W. and J.S. with contributions from all authors.

## Data and materials availability

Coordinates, light amplitudes, dark amplitudes and extrapolated structure factors for light activated data have been deposited in the PDB database under accession codes xxxx (TR-SMX M-state), xxx (TR-SMX N-state), xxxx (steady-state SMX M/N-state). Coordinates and dark state structure factors (steady-state SMX dark) have been deposited in the PDB database under accession code xxxx.

