## Supplementary Materials for "Proton uptake mechanism in bacteriorhodopsin captured by serial synchrotron crystallography"

**Supplementary Material**

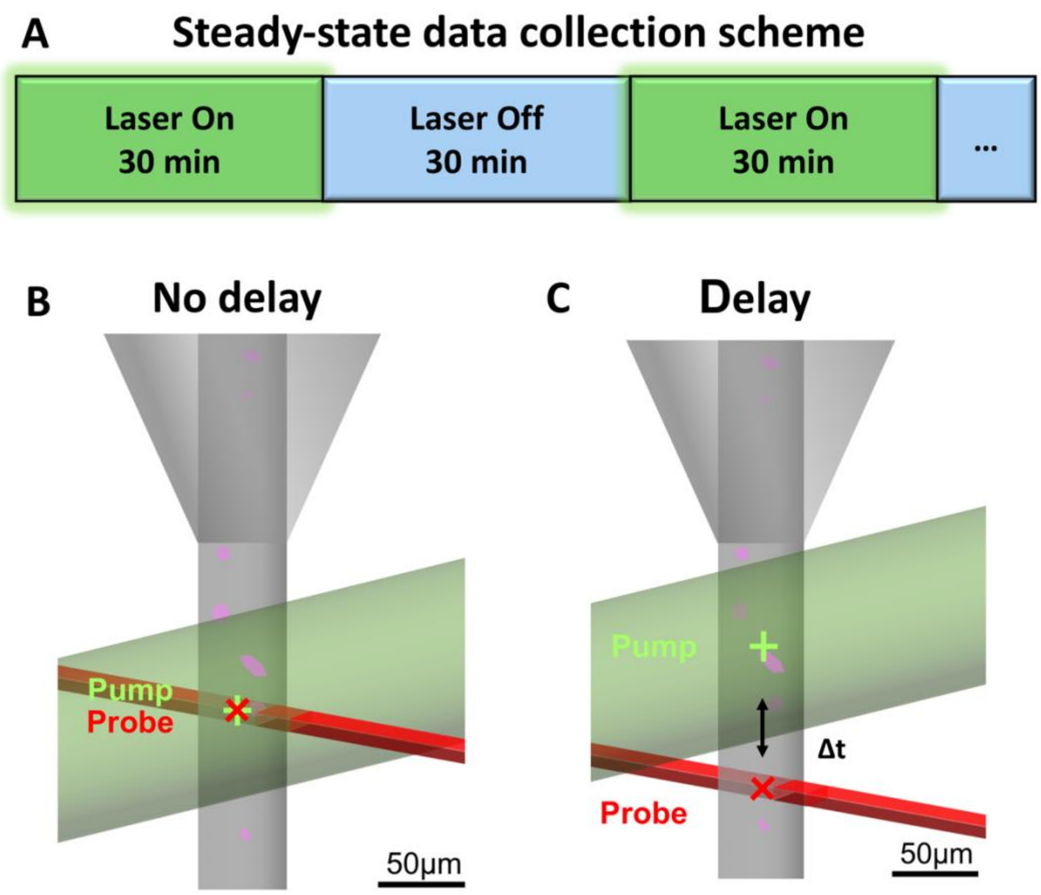


**Figure S1.** Steady-state-SMX data collection scheme. **(A)** Data are collected by interleaving data collection runs of 30 minutes with the laser on (green bars) and runs where the laser is switched off (blue bars). **(B)** Intersection scheme between X-rays and laser beam for steady-state-SMX to resolve dominant intermediates in a photocycle. **(C)** Intersection scheme between X-rays and the laser beam for very simple time-resolved experiments using a laser diode. The distance of the X-ray interaction region from the laser interaction region in combination with the jet speed determines the time delay between pump and probe.

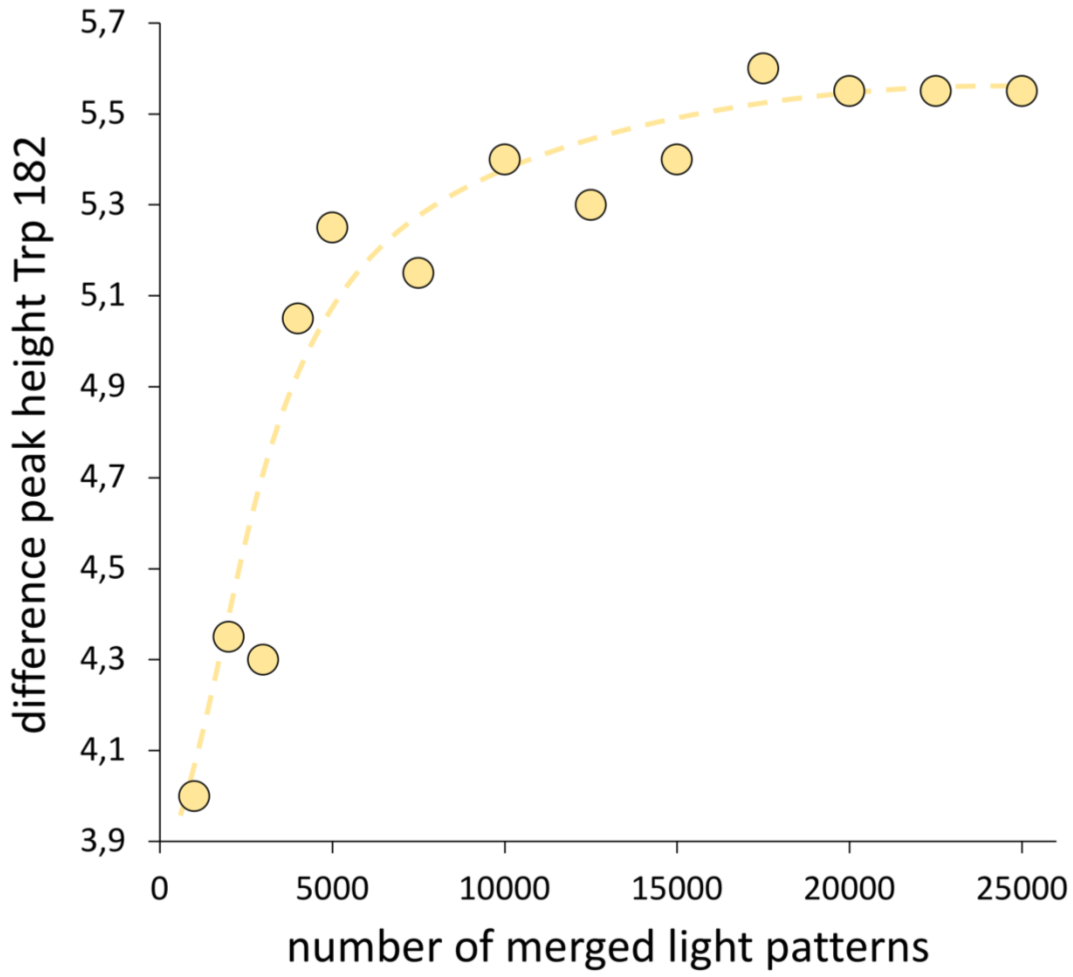


**Figure S2. Estimation for the number of diffraction images needed for a structural snapshot.** The quality of $F_o$(light)-$F_o$(dark) difference maps obtained from a time-resolved serial crystallography experiment increases with the number of included diffraction images because changes in the overall structure factor amplitudes are small and thus have to be measured accurately. To estimate the number of images needed in our experimental setup we calculated maps with defined subsets of the steady-state SMX data and compared the height of characteristic difference peaks. The graph plots the average positive and negative difference peaks around Trp182 against the number of light images (yellow circles, trend indicated as a dashed line) used to calculate the map. A plateau starting from about 10’000 images indicates convergence of data.

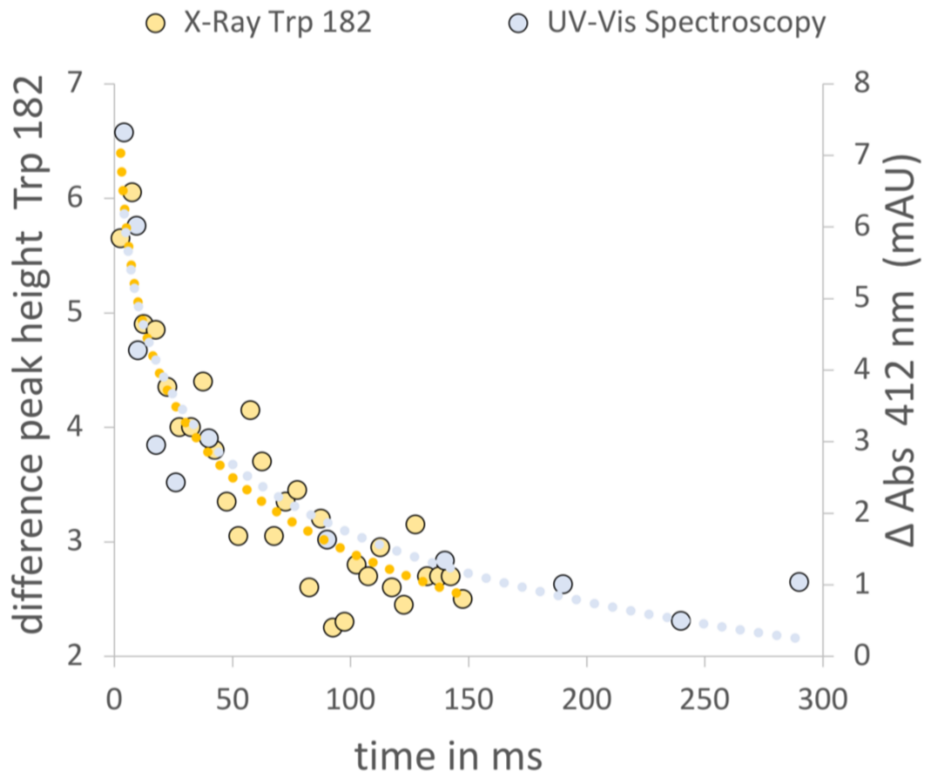


**Figure S3. Comparing the decay of late photointermediates using time-resolved absorption spectroscopy and crystallography.** Time-resolved absorption spectroscopy allows following photocycle kinetics using absorption changes at 412 nm (right axis, cyan circles). The plot compares spectroscopic data taken from Nogly *et al*. (Nogly et al. 2016) to the decay of late photo intermediates measured by the averaged difference peak height of Trp182 (left axis, yellow circles).

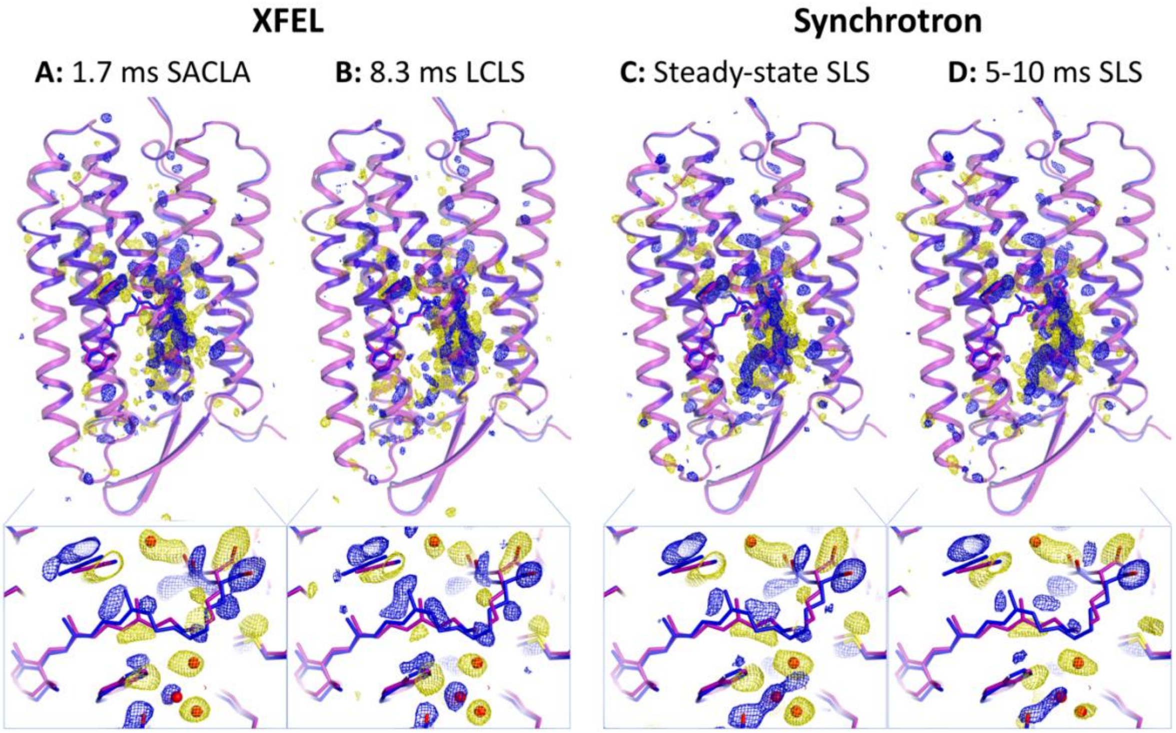


**Figure S4. Comparison of difference density maps obtained at XFELs and the synchrotron.** Overview of electron difference maps ($F_o$(light)-$F_o$(dark), negative (yellow) and positive (blue) contoured at 3 σ) collected (**A**) 1.7 ms after activation (data taken from (Nango et al. 2016)) at the Spring-8 Ångstrom Compact Free Electron Laser (SACLA), (**B**) 8.3 ms after activation (data taken from (Nogly et al. 2018)) at the Linac Coherent Light Source (LCLS) (**C**) with steady-state data and (**D**) 5-10 ms after activation at the Swiss Light Source (SLS). For easier comparison, all difference maps were calculated with the same settings and a resolution cut-off of 2.1 Å. As a quantitative measure of map quality, we have compared the positive peak of Trp182 (small insets showing dark-state in purple and M-state in blue) which indicates a shift of this residue once retinal reaches the planar 13-*cis* configuration. It is one of the strongest features in each map and reads out as 7.1 σ, 7.5 σ, 8.5 σ, and 8.3 σ in case of TR-SFX (1.7 ms, SACLA), TR-SFX (8.33 ms, LCLS), steady-state-SMX (SLS) and TR-SMX (5-10 ms, SLS), respectively. The quality of difference maps is determined by many interdependent factors including the number of collected images, the levels of activation, the achieved resolution but also fluctuations in beam quality and detector noise. Our direct comparison shows that despite these differences, a serial synchrotron experiment can provide difference maps of the same quality as those obtained from XFEL experiments but with much lower sample consumption and cost.

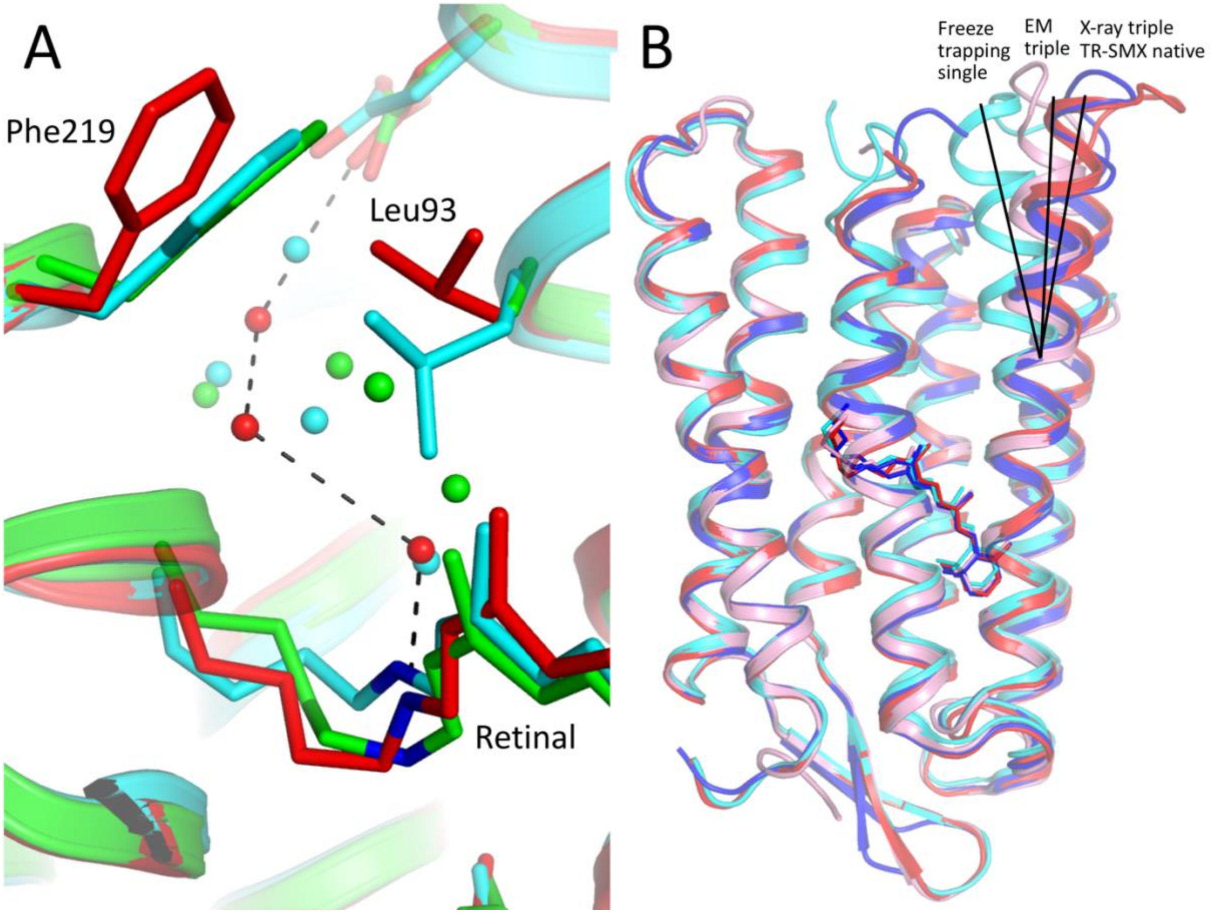


**Figure S5. Comparison of the N intermediate from TR-SMX with structures of bR mutants (A)** Overlay of the 10-15 ms dominating N-state (red), with the crystal structure obtained by cryo-trapping of the L93A mutant (green, pdb code: 3VI0 (Zhang et al. 2012), and the crystal structure obtained by cryo-trapping of the V49A mutant (cyan, pdb code: 1P8U (Schobert et al. 2003)). Water molecules close to the Grotthuss proton wire (dotted lines) in our structure are highlighted. Water positions in 3VI0 are shifted and additional waters are present in the free space created by the L93A mutation. While the V49A mutation in 1P8U shows the same water position close to the Schiff base, the waters upstream are again displaced into the free space this mutation creates. The mutation of Val49 to Ala further affects the interaction with Leu93, leading to differences in Leu93 and Phe219 positions which in turn prevent formation of the open conformation in our native 10-15 ms structure. **(B)** Overlay of the N-state at 10-15 ms (red), with the crystal structure of an N'-state of the Val49Ala single mutant (cyan, pdb code: 1P8U (Schobert et al. 2003)), and the dark-state of the Asp96Gly/Phe171Cys/Phe219Leu triple mutant of bR determined by electron microscopy (pink, 1FBK (Subramanlam & Henderson 2000),) and X-ray crystallography (blue, 4FPD (Wang et al. 2013)). Black lines signify the large shift on the cytoplasmic side of helix F. The large-scale motion of helix F resembles the outward movement of transmembrane helix 6 in G protein-coupled receptors as the principal conformational change that opens these receptors for binding of intracellular signaling partners. Perhaps evolution has preserved not only the basic seven-transmembrane helix fold but also a certain set of functional rearrangements within this important class of membrane proteins.

| *Dataset* | **Steady-state dark** | **Steady-state light** | **TR-SMX dark** | **TR-SMX 0-5 ms** | **TR-SMX 10-15 ms** |
|---|---|---|---|---|---|
| ***Space group*** | $P6_3$ | $P6_3$ | $P6_3$ | $P6_3$ | $P6_3$ |
| ***Unit cell (a = b, c in Å, α = β, γ in °)*** | 62.0 110.3 90 120 | 62.0 110.3 90 120 | 62.0 110.3 90 120 | 62.0 110.3 90 120 | 62.0 110.3 90 120 |
| ***frame rate*** | *50* | *50* | 200 | 200 | 200 |
| ***Collected images*** | 1’220’047 | 1’344’261 | 457’360 | 91’472 | 91’472 |
| ***Indexed patterns*** | 119’374 | 114’052 | 68’698 | 13’346 | 13’384 |
| ***patterns indexed (%)*** | 9.8 | 8.5 | 15.0 | 14.6 | 14.6 |
| ***Resolution*** | 30.78 – 1.8 (1.85 – 1.8) | 30.78 – 1.8 (1.85 – 1.8) | 38.45 -2.1 (2.18 – 2.1) | 38.45 -2.3 (2.37 – 2.3) | 38.45 -2.3 (2.37 – 2.3) |
| ***Number of Reflections*** | 34’896’314 | 30’552’664 | 13’671’870 | 2’726’066 | 2’699’539 |
| ***Number of unique Reflections*** | 21’977 | 21’976 | 13’882 | 10’589 | 10’589 |
| ***Multiplicity*** | 1’587.9 (198.2) | 1’390.3 (58.1) | 984.9 (24.6) | 257.4 (70.5) | 254.9 (69.8) |
| ***Completeness*** | 100 (100) | 100 (99.5) | 100 (99.9) | 100 (100) | 100 (100) |
| ***I / sigma*** | 14.3 (1.17) | 12.4 (0.36) | 13.3 (1.33) | 6.95 (0.92) | 6.77 (0.94) |
| ***CC**** | 0.99 (0.69) | 0.99 (0.31) | 0.99 (0.83) | 0.99 (0.32) | 0.99 (0.49) |
| ***$CC^{1/2}$*** | 0.99 (0.31) | 0.99 (0.05) | 0.99 (0.52) | 0.99 (0.06) | 0.99 (0.13) |
| ***Rsplit*** | 2.6 (106.1) | 2.4 (403.0) | 3.4 (85.7) | 7.4 (153.7) | 7.5 (173.0) |
| ***Activation level*** | - | 0.38 | - | 0.24 | 0.18 |

**Table S1. Data Statistics**

| *Dataset* | **Steady-state dark** | **Steady-state M-/N-states** | **TR-SMX M-state** | **TR-SMX N-state** |
|---|---|---|---|---|
| ***Space group*** | *P*6$_3$ | *P*6$_3$ | *P*6$_3$ | *P*6$_3$ |
| ***Unit cell (a = b, c in Å, α = β γ in °)*** | 62.0 110.3 90 120 | 62.0 110.3 90 120 | 62.0 110.3 90 120 | 62.0 110.3 90 120 |
| ***Resolution*** | 30.3 - 1.8 (1.86 - 1.8) | 30.3 - 2.0# (2.1 - 2.0) | 38.4 - 2.6# (2.98 - 2.60) | 31.0 - 2.6# (2.98 - 2.60) |
| ***Number of unique Reflections*** | 22272 (2230) | 15903 (2127) | 6737 (1926) | 6474 (1838) |
| ***Completeness*** | 100 (100) | 98.1 (99) | 90.1 (84) | 87.3 (80) |
| ***Datasets*** | steady-state dark | steady-state light/dark | TR-SMX 0-5 ms/dark | TR-SMX 10-15 ms/dark |
| ***Activation level*** | - | 0.38 | 0.24 | 0.18 |
| ***Occupancy M / N*** | - | 60/40 | 100/0 * | 0 /100 * |
| ***$R_{cryst}$ / $R_{free}$*** | 17.4 (19.8) | 20.1 / 24.6 (31.1 / 35.2) | 24.9 / 30.1 (37.2 / 47.5) | 31.7 / 36.0 (47.2 / 54.5) |
| ***Ramachandran favored / allowed / outliers*** | 220 / 6 / 1 | 418 / 20 / 4 | 220 / 6 / 1 | 215 / 6 / 0 |
| ***RMS bonds*** | 0.018 | 0.007 | 0.003 | 0.002 |
| ***RMS angles*** | 1.23 | 0.87 | 0.45 | 0.44 |
| ***Average B-factor*** | 74.0 | 44.0 | 83.0 | 92.0 |
| ***PDB Code*** | XXXX | XXXX | XXXX | XXXX |

**Table S2. Refinement Statistics**

#Light data extrapolation lowers the data resolution usable for refinement depending on the activation level but allows to selectively analyze the activated state. *Timepoint 0-5 ms contains a fraction of N-state and timepoint 10-15 ms contains a small fraction of M-state (see Figure 2C). Here, we refined only the predominant model against extrapolated data.

**Movie S1: Evolution of difference density over time.** A three-dimensional movie of the rise and decay of overall structural changes. The structures of the dark (purple), M-state (blue) and N-state (red) are shown as a cartoon with retinal as sticks. The evolution of difference electron density (negative (yellow) and positive (blue) contoured at 3 σ) is shown for the first 20 frames during which bR completes its photocycle.

**Movie S2: Dynamic view of the proton uptake mechanism.** The movie shows the transition between three structural intermediates: dark state, M-state and N-state. The morphs between the structures were generated with Chimera and highlight the main structural arrangements on the cytoplasmic site and formation of the water chain from the proton donor Asp96 towards the Schiff base.
